# Elucidating the H^+^ coupled Zn^2+^ transport mechanism of ZIP4; implications in *Acrodermatitis Enteropathica*

**DOI:** 10.1101/474528

**Authors:** Eitan Hoch, Israel Sekler

## Abstract

Cellular Zn^2+^ homeostasis is tightly regulated and primarily mediated by designated Zn^2+^ transport proteins, namely ZnTs (SLC30) that shuttle Zn^2+^ efflux, and ZIPs (SLC39) that mediate Zn^2+^ influx. While the functional determinants of ZnT-mediated Zn^2+^ efflux are elucidated, those of ZIP transporters are lesser understood. Previous work has suggested three distinct molecular mechanisms: (I) HCO3^−^ or (II) H^+^ coupled Zn^2+^ transport, or (III) a pH regulated electrodiffusional mode of transport. Here, using live-cell fluorescent imaging of Zn^2+^ and H^+^, in cells expressing ZIP4, we set out to interrogate its function. Intracellular pH changes or the presence of HCO3^−^ failed to induce Zn^2+^ influx. In contrast, extracellular acidification stimulated ZIP4 dependent Zn^2+^ uptake. Furthermore, Zn^2+^ uptake was coupled to enhanced H^+^ influx in cells expressing ZIP4, thus indicating that ZIP4 is not acting as a pH regulated channel but rather as an H^+^ powered Zn^2+^ co-transporter. We further illustrate how this functional mechanism is affected by genetic variants in *SLC39A4* that in turn lead to *Acrodermatitis Enteropathica*, a rare condition of Zn^2+^ deficiency.

## Introduction

Zn^2+^ is an essential nutrient that plays key roles in a variety of cellular and physiological processes ^1^. It is therefore not surprising that Zn^2+^ deficiency, underlined by an inability to acquire nutritional Zn^2+^, has devastating effects. These range from mental disorders, to Immune system dysfunction and growth retardation ^2^. The importance of Zn^2+^ to human physiology is further emphasized by a recent finding that approximately 2800 proteins (~10% of the human proteome) are potentially Zn^2+^ binding; these include transcription factors, Zn^2+^ finger proteins, and a variety of enzymes ^3^. Yet, little is known about the process of Zn^2+^ uptake and how Zn^2+^ ions move across membranes and into cells and organelles.

Two groups of mammalian Zn^2+^ transporters have been identified; SLC39 (ZIP; ZRT-IRT-like protein) mediate Zn^2+^ influx, and SLC30 (ZnT; Zinc transporters) mediate Zn^2+^ efflux ^4^.

The 10 members of the ZnT family of efflux transporters have been linked to numerous cellular processes that include insulin secretion ^5, 6^ and TNAP activation ^7, 8^. The functional mechanism of these transporters has been studied in a variety of models, from human cell cultures ^9^ to plants ^10^ and bacteria ^11^; all indicating a Zn^2+^/H^+^ exchange mechanism. The recently solved structure of a bacterial ZnT orthologue ^12^ has further enhanced our knowledge on the biochemical and biophysical properties of this group.

The 14 members of the ZIP family mediate transport of Zn^2+^ ions into the cytoplasm, either from the extracellular surroundings of the cell, or from intracellular organelles ^13^. Members of this group have been linked to various pathologies, such as Ehlers-Danlos syndrome ^14, 15^, and cadmium toxicity ^16^. In contrast to ZnTs, our understanding of the mechanisms that govern Zn^2+^ transport by this group is lacking.

ZIPs typically have 8 TMDs, with both N- and C- termini facing the extracytoplasmic side, and a Histidine rich domain is found in the cytoplasmic loop between TMDs 3 and 4. The role of this loop is undetermined; however, mutating these residues in the Yeast orthologue Zrt1 resulted in different localization of the protein, with no effect on Zn^2+^ transport ^17^. TMDs 4 and 5 are conserved ^18^ and highly amphipathic, and thus have been suggested to form a cavity through which ion transport is mediated ^4^. Molecular modeling of ZIP4 (Antala, 2015) has recently supported this, and further experimental corroboration comes from IRT1, from *Arabidopsis Thaliana*, in which mutating charged residues in TMDs 4 and 5 reduced Fe^2+^ uptake, and reciprocally increased Zn^2+^ uptake ^19^. Interestingly, mutating a charged Histidine residue in the catalytic core of ZnTs, alters Zn^2+^ vs. Cd^2+^ selectivity ^20^.

In the current report, we focus on ZIP4 that plays an important role in acquiring nutritional Zn^2+ 21^. ZIP4 is highly expressed in the small intestines and the embryonic visceral yolk sac, where it primarily localizes to the apical PM, and undergoes rapid endocytosis, following exposure to Zn^2+ 22, 23^. Under conditions of Zn^2+^ deficiency, ZIP4 is apparently cleaved and a shorter peptide of 37-40 kDa is detected at the PM ^21, 23, 24^, suggesting proteolytic processing regulates ZIP4 expression.

The importance of this transporter is emphasized in individuals with Acrodermatitis Enteropathica (AE), a rare human genetic disorder. AE is manifested by several variants of the SLC39A4 gene ^25-28^ that lead to Zn^2+^ deficiency, characterized by skin lesions, growth retardation, immune system dysfunction and neurological disorders ^2, 29^. The 3D-structure of BbZIP, a prokaryotic ortholog, was recently identified and several AE-associated variants were mapped onto a ZIP4 model that was based on the solved structure. These variants are clustered around the transmembrane ZIP4 domains and are thought to be critical for ZIP4 homodimerization ^30^. ZIP4 has also been signified as a marker for pancreatic cancer ^31^, leading to elevated Zn^2+^ content in tumor cells, and thus increased cell proliferation and tumor size. Reciprocally, ZIP4 down regulation had a protective effect, limiting tumor growth ^32^. Despite the importance of this transporter to human health, the molecular mechanisms by which it mediates Zn^2+^ uptake are unknown.

Previous studies performed on mammalian members of the ZIP family have suggested Zn^2+^ uptake is enhanced either under alkaline conditions or following the addition of HCO3^−^, thus suggesting a Zn^2+^/HCO3^−^ co-transport mechanism. This was suggested for ZIP2 ^33^, ZIP8 ^34^ and ZIP14 ^35^. On the contrary, studies performed on FrZIP2, a close homologue to ZIP3, obtained from *Takifugu rubripes* (Puffer fish) have shown a reduction of Zn^2+^ uptake following the addition of HCO3^−^ and suggested an increase of Zn^2+^ uptake under acidic pH conditions, suggesting a possible Zn^2+^/H^+^ co-transport mechanism ^36^. A recent study has mapped the catalytic core of ZIP4 suggesting a pentahedral Zn^2+^ coordination site composed of 3 Histidine and 2 aspartate residues ^37^. Furthermore, recent studies performed on a purified and reconstituted ZIP bacterial homologue, ZIPB, suggest it acts as a pH regulated slow electrodiffusional channel, and not a transporter, mediating Zn^2+^ transport that is uncoupled from HCO3^−^ or H^+^ transport ^38^.

Here we monitor cytoplasmic Zn^2+^ and pH changes in HEK293-T cells. Our results indicate that in contrast to the channel-like behavior of the bacterial transporter, in ZIP4 transport of Zn^2+^ and H^+^ are coupled, supporting a Zn^2+^/H^+^ co-transport mode. This suggests that ZIP4 has undergone an evolutionary transformation form channel to transporter. We further provide a functional basis for two SLC394 genetic variants linked to Zn^2+^ deficiency in AE patients.

## Results

### Zn^2+^ transport by ZIP4

Previous studies have shown that ZIP4, as well as other members of the ZIP family, undergoes rapid endocytosis in the presence of extracellular Zn^2+ 22, 23^, thus constituting a major experimental challenge in directly monitoring the transport mechanism of ZIP4.

Therefore, we initially asked if the rates of transport and endocytosis are sufficiently different to distinguish between. The rate of endocytosis was monitored using the well-established ZIP4 surface-labeling protocol ^22^.

Zn^2+^ (50uM) was added to HEK293-T cells expressing HA-tagged mZIP4 as indicated (Fig. 1A). Cells were then washed with ice-cold PBS, and immediately transferred to ice, in order to stop any endocytosis processes. Following, cells were fixed in PFA and exposed to anti-HA antibodies. Unbound antibodies were extensively washed and bound HA was determined as a function of surface ZIP4 expression by WB analysis of the anti-HA tag antibody.

**Figure 1:**
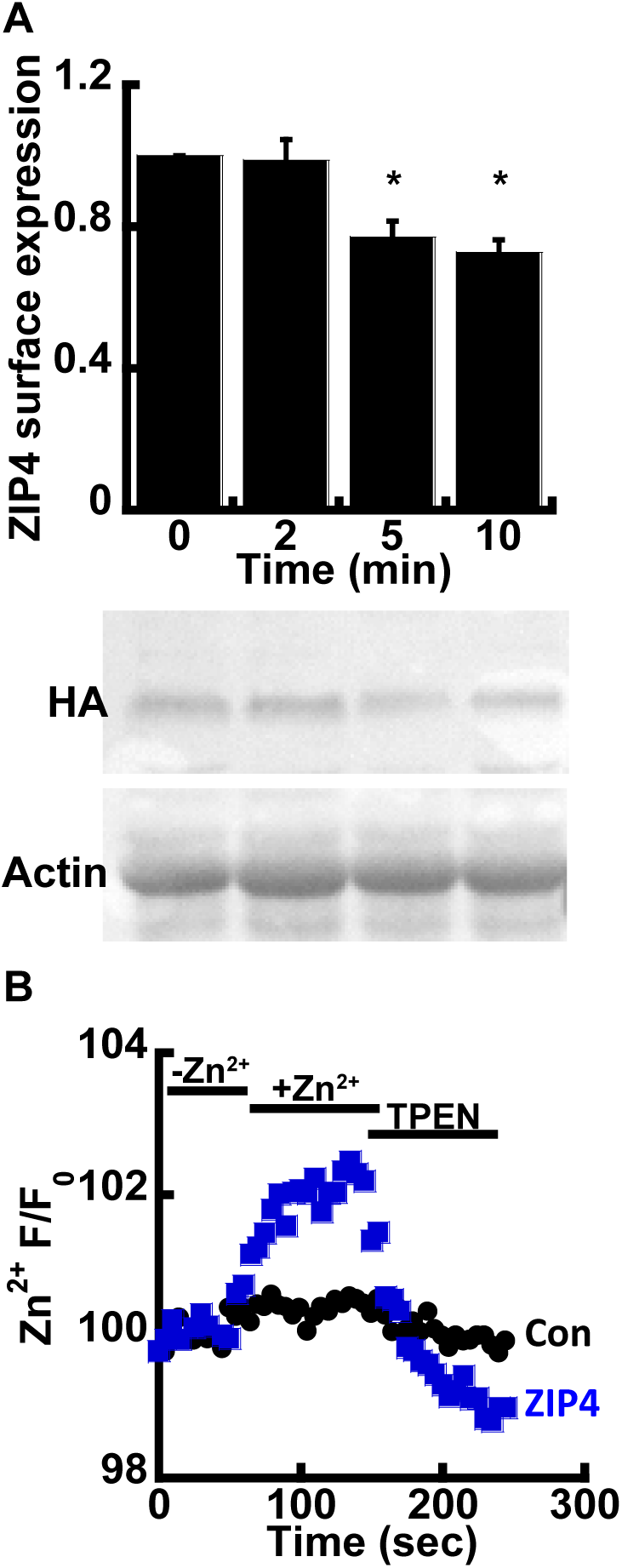
ZIP4 Zn^2+^ transport assay. **(A)** Immunoblot analysis and normalized surface expression of HEK-293T cells transfected with HA-tagged mZIP4 and exposed to 50µM Zn^2+^, as indicated. **(B)** Experimental assay used to monitor Zn^2+^ uptake in HEK-293T cells transfected with an empty control vector (black) and mZIP4 (blue). Cells were loaded with 1µM Fluozin-3AM and subjected to 50µM Zn^2+^ and 50µM TPEN, as indicated.

Consistent with pervious results, no internalization of ZIP4 was observed during the first 2 minutes of Zn^2+^ exposure (Fig. 1A) and a reduction in ZIP4 surface expression was only monitored after 5 minutes. Our ensuing transport assays were therefore set to a 2-minute time interval, following the addition of Zn^2+^, thus allowing accurate monitoring of Zn^2+^ transport, uninterrupted by ZIP4 endocytosis.

Zn^2+^ transport by ZIP4 was monitored in HEK293-T cells overexpressing mZIP4 and preloaded with 1uM Fluozin-3AM, a Zn^2+^ sensitive fluorescent probe, commonly used for monitoring Zn^2+^ transport ^20, 39^. Cells were perfused in Ringer’s solution containing 50uM Zn^2+^ and the rate of Zn^2+^ transport was measured and compared to cells transfected with a control vector. Zn^2+^ transport rates mediated by ZIP4 expressing cells were ~5 fold higher than control cells (Fig. 1B) indicating that the expression of ZIP4 is linked to enhanced Zn^2+^ transport. To ascertain the increase in Fluozin-3AM fluorescence is triggered by cytoplasmic Zn^2+^, the Zn^2+^ sensitive intracellular chelator TPEN was added at the end of the experiment, following which cytoplasmic fluorescence returned to baseline levels, thus indicating that the fluorescent signal is mediated by changes in cytosolic Zn^+^. Altogether the results of this part indicate that expression of ZIP4 leads to enhanced Zn^2+^ influx across the PM.

### Zn^2+^ uptake by ZIP4 is pH dependent

We next sought to determine the mechanism that drives Zn^2+^ transport by ZIP4. In other members of the ZIP family, Zn^2+^ transport was suggested to be coupled to HCO3^−33-35^, and we therefore sought to determine the effect of HCO3^−^ on Zn^2+^ transport mediated by ZIP4. To address this, HEK293-T cells were transfected with either ZIP4 or an empty control vector, and Zn^2+^ transport was compared in cells perfused with pH7.4 Ringer’s solution containing 50uM Zn^2+^, in the presence or absence of 20uM NaHCO3 (Figure 2A). No significant differences were observed, and our results, therefore, do not support a Zn^2+^/HCO3^−^ coupled transport mechanism for ZIP4.

**Figure 2:**
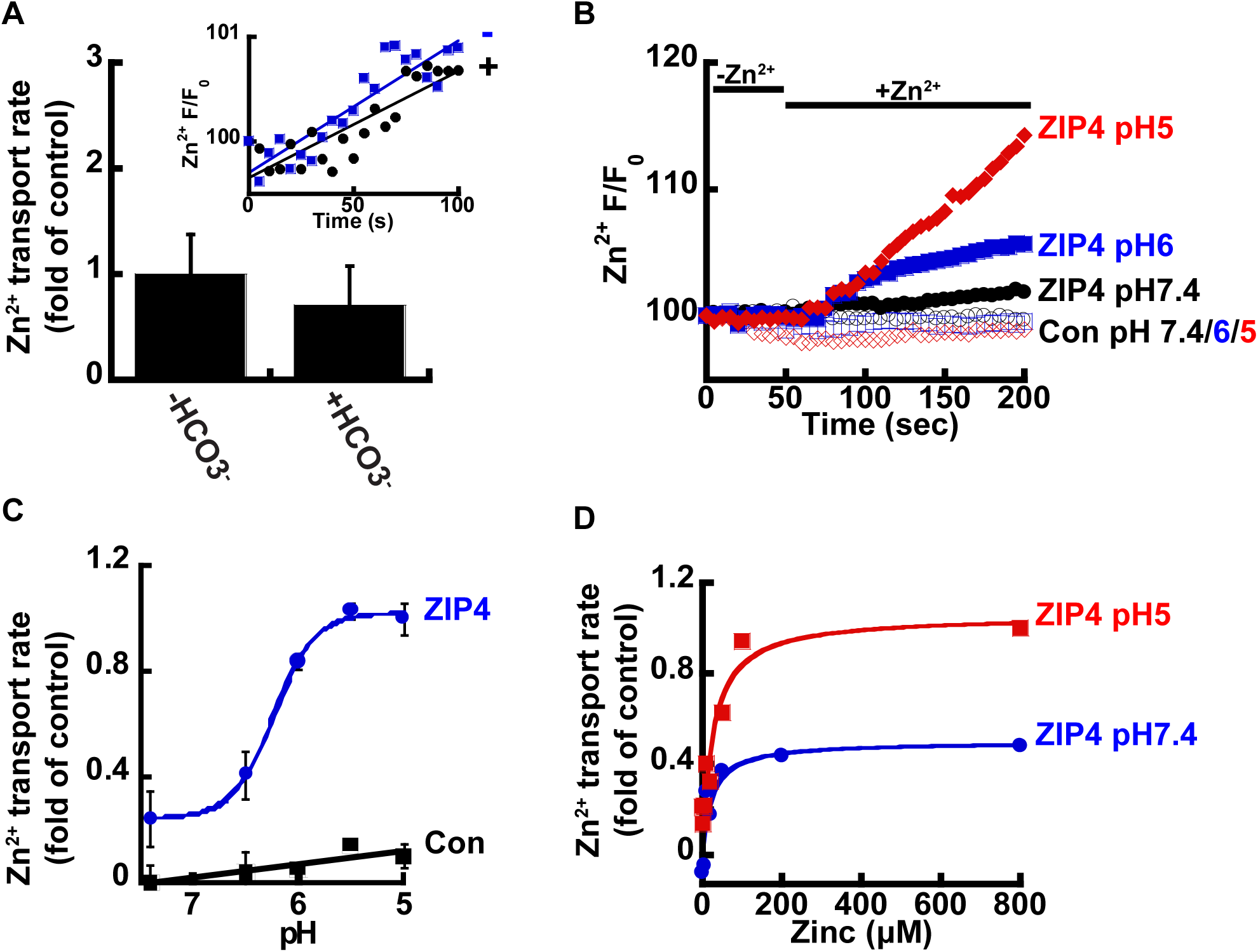
ZIP4-mediated Zn^2+^ transport is pH dependent. (A) HCO3-has no effect on Zn2+ uptake. HEK-293T cells transfected with ZIP4 were loaded with 1µM Fluozin-3AM and loaded with 50µM Zn^2+^ in HEPES buffered Ringer’s solution (grey) or NaHco3^−^ buffered Ringer’s solution (black). Zn^2+^ uptake was normalized and compared to controls. (B) Representative traces of Zn^2+^ uptake, in HEK293-T cells, transfected with an empty control vector (empty symbols) or mZIP4 (full symbols) under varying pH levels, as indicated. (C) Normalized rates of Zn^2+^ uptake by cells expressing either mZIP4 or a control vector. (D) Zn^2+^ uptake, HEK293-T cells expressing ZIP4, at pH7.4 (blue) and pH5 (red), at the indicated zn^2+^ concentrations (0-800µM).

Studies performed on ZIP homologues from other species, such as the bacterial ZIPB ^38^ and puffer fish FrZIP2 ^36^, suggest ZIPs act as H^+^ activated Zn^2+^ channels that are independent of HCO3-. To determine the effect of pH on Zn^2+^ transport, by ZIP4, we monitored Zn^2+^ transport at the indicated pH values (Fig. 2B). Zn^2+^ transport shows strong pH dependency, with a 4-fold increase of ZIP4 mediated Zn^2+^ transport rates at pH5 compared to pH7.4 (Fig. 2C). In contrast, no increase in Zn^2+^ transport was monitored in control cells at pH5, when compared to pH7.4, indicating that this pH effect is related to ZIP4 activity and not to a non-selective pH dependent change in Zn^2+^ concentrations that may occur by cytosolic acidification ^40^. Sensitivity of Zn^2+^ transport to pH was similar to that documented for the bacterial and Puffer fish homologues ^36, 38^. Hill’s coefficient was calculated at 5.08 ±0.8, indicating a Zn^2+^/H^+^ stoichiometry of 1:5.

To determine if pH controls the apparent affinity or rate of ZIP4 mediated Zn^2+^ transport, we conducted Zn^2+^ dose-response experiments at pH5 and pH7.4, utilizing the same experimental design described previously, with Zn^2+^ concentrations ranging from 0-800uM (Fig. 2D). The results were fitted to a Michaelis-Menten equation and are summarized in Table. 1. While the affinity for Zn^2+^ transport (Km) was pH independent, the rate of ion transport (Vmax) doubled from 0.49796s^−1^ at pH7.4 to 1.0554s^−1^ at pH5, indicating that acidic pH accelerates the turnover rate of Zn^2+^ transport, with no effect on affinity.

Acidic extracellular pH will lead to an intracellular pH drop. The latter can in turn trigger an intracellular Zn^2+^ rise that is independent of ZIP4 activity, by enhancing the dissociation of intracellular bound Zn^2+ 39, 40^. In addition, the observed activation of ZIP4 may not necessarily be through an extracellular effect, but by a change in cytosolic pH. To address this possibility, we applied the well documented ammonium pre-pulse paradigm ^41^ that selectively triggers an intracellular, but not extracellular, pH change, in cells preloaded with either 1uM BCECF-AM (Fig.3A) or 1uM Fluozin-3AM (Fig.3B). All solutions were Na^+^ free, to prevent the activation of H^+^ efflux via the major cytosolic acid extruder, the Na^+^/H^+^ exchanger.

**Figure 3:**
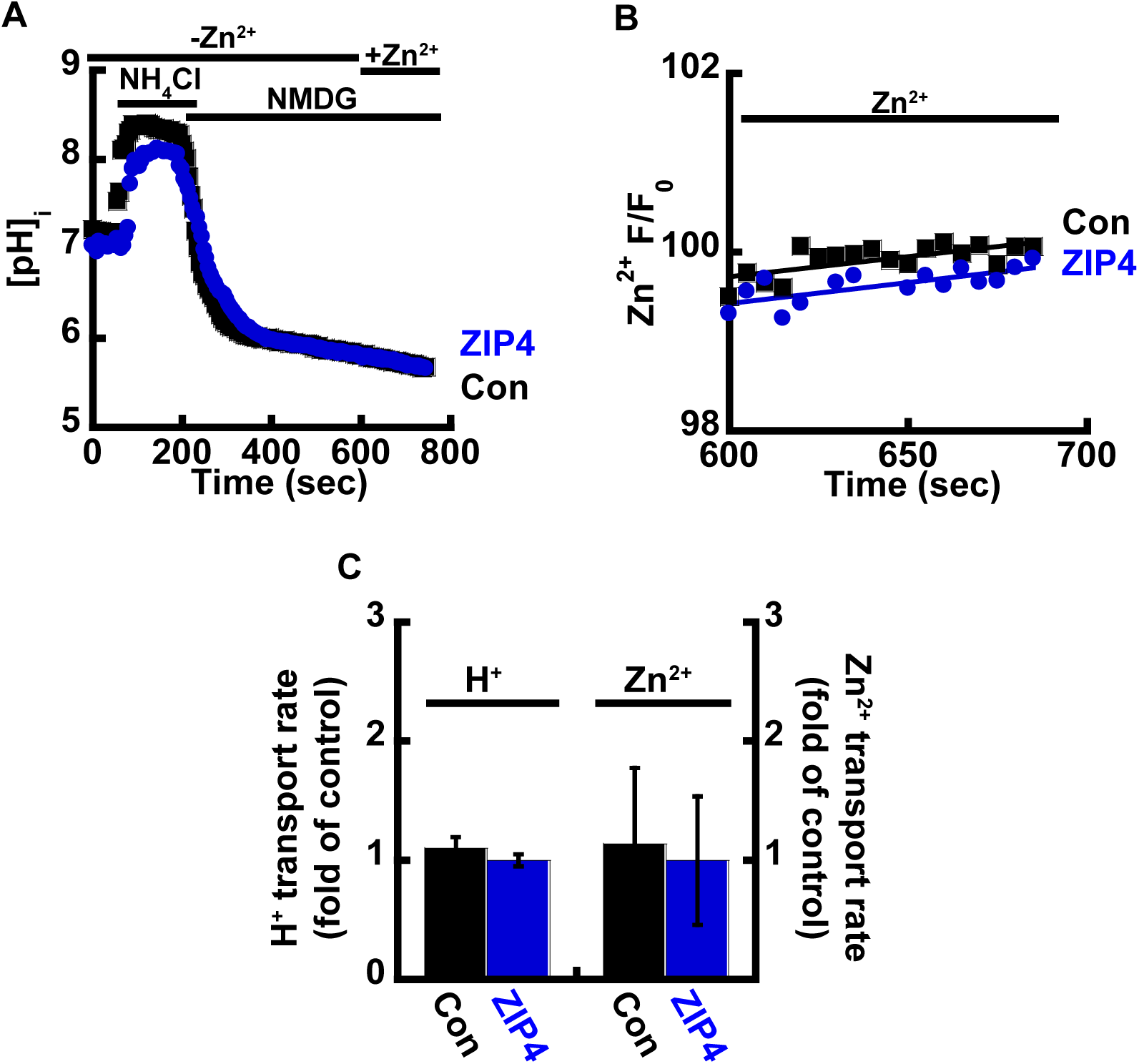
Intracellular acidification limits Zn^2+^ uptake. **(A)** Representative traces of HEK293-T cells transfected with ZIP4 (blue) or an empty control vector (black), loaded with 1µM BCECF-AM, and subjected to ammonium pre-pulse paradigm to monitor cytoplasmic pH changes without extracellular acidification. **(B)** Corresponding Zn^2+^ traces in cells loaded with 1µM FLuozin-3AM. **(C)** Normalized H^+^ and Zn^2+^ transport rates.

Following intracellular acidification, 50uM Zn^2+^ were added and both Zn^2+^ and H^+^ transport rates were compared in Zip4 or control cells (Fig.3C). No significant differences were observed indicating the pH activation of Zn^2+^ transport relies solely on extracellular protons. Furthermore, intracellular acidification triggered an inhibition of Zn^2+^ transport by ZIP4 indicating that the reversal of the H^+^ gradient potentially blocked Zn^2+^ transport. Note that a similar result was obtained with FrZIP2, in which Zn^2+^ transport was inhibited following the addition of extracellular HCO3^−36^.

### Zn^2+^ and H^+^ transport by ZIP4 are coupled

The above results strengthen the hypothesis that extracellular H^+^ ions generate the driving force for ZIP4, suggesting two possible modes of transport for ZIP4: (1) H^+^/Zn^2+^ co-transporter and (2) H^+^ sensitive Zn^2+^ channel. To distinguish between these modes of operation, H^+^ transport was monitored in ZIP4 expressing cells preloaded with 1uM BCECF-AM. We reasoned that if ZIP4 acts as a channel no ZIP4 mediated proton transport will be observed.

In the absence of Zn^2+^, no differences in H^+^ transport were observed between ZIP4 expressing cells and controls (Fig. S1A, S1B). In the presence of Zn^2+^, at neutral pH, no differences were observed either (Fig. 4A). Note that at neutral pH in the presence of Zn^2+^, H^+^ flux was slightly reduced (Fig. S1C). This is consistent with the blocking effect of Zn^2+^ on multiple cationic channels that are permeable to H^+^. In the presence of Zn^2+^, at acidic pH, a clear rise in cell acidification was observed in ZIP4 expressing cells, when compared to control cells (Fig. 4A, 4B).

**Figure 4:**
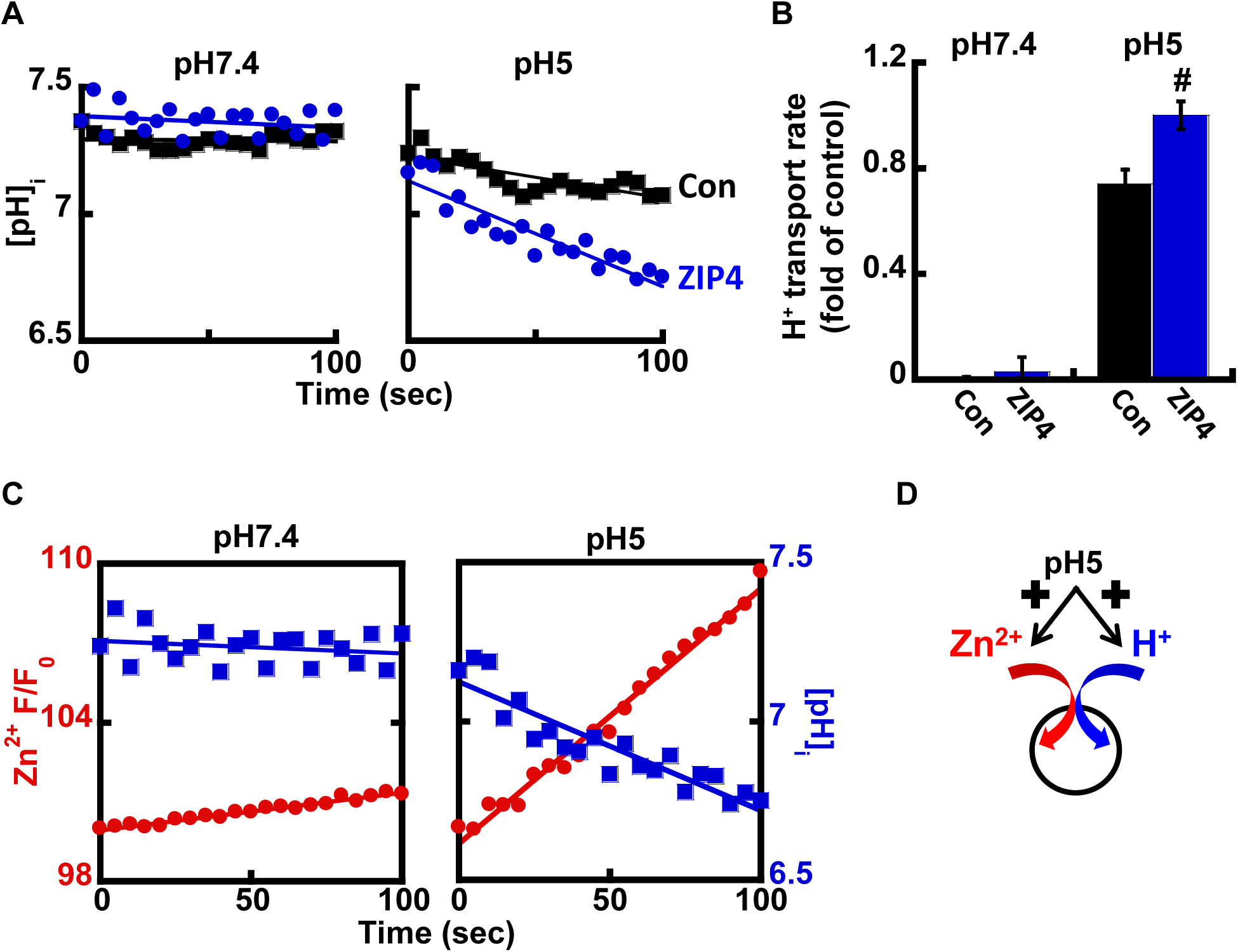
ZIP4 mediates H^+^ coupled Zn^2+^ transport. (A) Representative traces of HEK293-T cells transfected with ZIP4 (blue) or a control vector (black), loaded with 1µM BCECF-AM, and monitored for cytoplasmic pH changes at pH7.4 (left) or pH5 (right), following the addition of 50µM Zn^2+^. **(B)** Normalized H^+^ transport rates recorded from control (black) or ZIP4 (blue) expressing cells. **(C)** Representative traces of Zn^2+^ (red) and H^+^ transport (blue), recorded with Fluozin-3AM and BCECF-AM accordingly. Note that H^+^ uptake is parallel to Zn^2+^ uptake. **(D)** Illustration of the suggested mechanisms of ZIP4.

Thus, our results show a robust coupling between Zn^+^ and H^+^ transport. Further support for a H^+^/Zn^2+^ co-transport mechanism is presented in figure 4C that compare the rate of Zn^2+^ transport and pH changes in cells expressing ZIP4, at pH7.4 (left panel) vs. pH5 (right panel). Note the strong reciprocity between cytosolic pH acidification and Zn^2+^ influx, which are both enhanced at acidic pH. Altogether our results suggest that ZIP4 mediates H^+^/Zn^2+^ co-transport. Under neutral pH conditions, both Zn^2+^ and H^+^ fluxes are subtle. In an acidic extracellular environment, Zn^2+^ and H^+^ influx rates are strongly increased, supporting a H^+^/Zn^2+^ co-transport mechanism (Fig. 4D).

### Acrodermatitis Enteropathica associated variants disrupt Zn^2+^ transport by ZIP4

Genetic variants in SLC39A4 are linked to Zn^2+^ deficiency in humans ^25-28^, but zinc supplementation has not always proven a useful treatment, implying different molecular mechanisms originating from different variants. Several SLC39A4 coding variants were previously tested and displayed varying levels of expression, as well as varying levels of Zn^2+^ uptake ^28^.

We focused on two variants of the SLC39A4 gene, for which surface expression was previously reported ^28^ and reproduced in our hands (Fig. 5A). The P200L variant is situated at the cytoplasmic N-terminus domain, and G539R within the loop connecting TMDs 4 and 5 that form the catalytic site (Fig. 5B).

**Figure 5:**
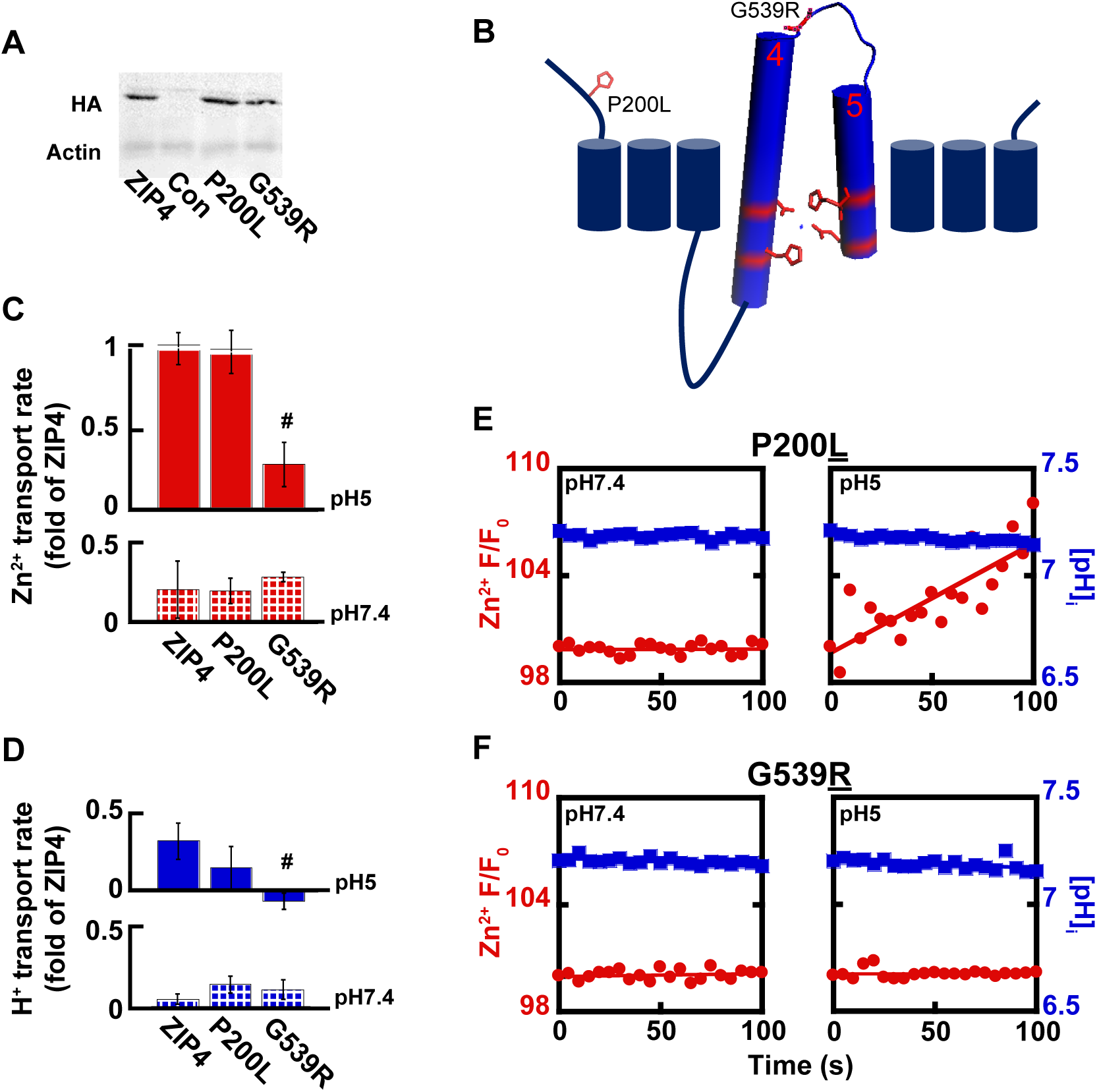
Genetic variation in ZIP4 affects catalytic and non-catalytic domains. **(A)** Immunoblot analysis of AE-associated ZIP4 constructs, as indicated, from HEK293-T cell lysates. **(B)** Membrane orientation of ZIP4 illustrates the position of AE-associated variants P200L and G539R. **(C)** Normalized rates of Zn^2+^ uptake at pH7.4 (bottom panel -striped) and pH5 (top panel - full), mediated by AE-associated ZIP4 mutants, in HEK-T cells loaded with Fluozin-3AM. **(D)** Normalized rates of H^+^ uptake at pH7.4 (bottom panel - striped) and pH5 (top panel - full), mediated by AE-associated ZIP4 mutants, in HEK-T cells loaded with BCECF-AM. Representative traces of Zn^2+^ (red) and H^+^ uptake (blue) recorded in HEK-T cells transfected with ZIP4^P200L^ **(E)** and ZIP4^G539R^ **(F)**.

Zn^2+^ and H^+^ transport, mediated by ZIP4^P200L^ were no different from those of the wild type ZIP4 (Fig. 5 C-E), suggesting that the P200L variant is not catalytically linked to the transport activity of ZIP4. Thus, Zn^2+^ deficiency observed in patients harboring this variant may be linked to other processes, e.g. cell surface dynamics of this transporter.

Zn^2+^ transport by ZIP4^G539R^, on the other hand, was not activated at acidic pH conditions and maintained basal activity at both pH7.4 and pH5 (Fig. 5C, 5D, 5F), suggesting a role for this residue in pH activation of ZIP4. A prediction for such a role is that H^+^ transport will also be affected. When assayed for H^+^ transport, cells expressing ZIP4^G539R^ demonstrated a 40% reduction at pH5, supporting a role of this residue in pH activation of the transporter, possibly due to its proximity to the catalytic core formed by TMDs 4 and 5. Thus, the reduction in both Zn^2+^ and H^+^ transport mediated by ZIP4^G539R^ suggests that the substitution of the small non-charged Glycine, in healthy individuals, to a larger charged Arginine, encountered in AE patients, disrupts the coupling of the H^+^ driving force and Zn^2+^ transport.

## Discussion

### ZIP4 mediates H^+^/Zn^+^ co-transport

ZIP4 is a membrane embedded protein, enriched in enterocytes, where it mediates Zn^2+^ uptake. Previous work has shown a regulatory process in which ZIP4 undergoes rapid endocytosis following the addition of extracellular Zn^2+ 22, 23^, thus constituting a major challenge in focusing directly on the transport mechanism of ZIP4. To overcome this difficulty, we monitored the timing of ZIP4 removal from the membrane. Endocytosis only begins 2 minutes following Zn^2+^ exposure. Using live cell imaging, we monitored direct ion transport by ZIP4 during this time thereby addressing the concern that ZIP4 surface expression is changing while its activity is assayed.

The mechanism by which ZIPs transport Zn^2+^ is by and large unclear. Studies of mammalian ZIPs ^33-35^ have suggested an HCO3^−^ dependent co-transport mechanism, with increased rates of transport either at alkaline pH or following the addition of HCO3^−^. Other studies on the bacterial ZIP homologue, ZIPB ^38^ indicated that it acts as a pH regulated, electrogenic facilitated diffusion channel, while studies on the puffer fish ZIP homologue, FrZIP2, indicated H^+^ Zn^2+^ co-transport.

Our data does not support the involvement of HCO3^−^. The addition of HCO3^−^ did not lead to elevated rates of Zn^2+^ transport, as previously reported also for FrZIP2 ^36^, and ZIPB ^38^.

Our results indicate that Zn^2+^ transport by mammalian ZIP4 is mediated by Zn^2+^/H^+^ co-transport, based on the following findings: 1) Extracellular acidification triggered ZIP4 dependent Zn^2+^ uptake. 2) In contrast, intracellular acidification - while extracellular pH was kept neutral - using the NH4Cl pre-pulse paradigm, inhibited ZIP4 dependent Zn^2+^ uptake. 3) Most relevant, ZIP4 dependent Zn^2+^ uptake - triggered by extracellular acidification - is linked to ZIP4 dependent intracellular acidification. These finding strongly suggest that ZIP4 functions as an H^+^/Zn^2+^ co-transporter.

We suggest that the mammalian ZIP4 transporter underwent an evolutionary progression from a facilitated diffusion channel to an H^+^ coupled Zn^2+^ transporter. What is the physiological advantage of H^+^/Zn^2+^ co-transport as opposed to facilitated diffusion? Facilitated diffusion mediators are built to support substrate gradient driven transport. Indeed, in bacteria, free intracellular Zn^2+^ is vanishingly low ^42^ and such a mechanism would therefore be optimal for Zn^2+^ uptake. In mammalian cells, on the other hand, Zn^2+^ is a rare but essential micronutrient that requires an optimal mechanism for maximal absorption.

A channel-like mode of transport is optimal for fast charge movements across the plasma membrane, but is less efficient for active transport against a concentration gradient, as harnessed by secondary active transporters. Zn^2+^ is an essential rate-limiting nutrient for eukaryotic cells, and Zn^2+^ deficiency is a frequent event with severe physiological consequences for many mammalian species. Thus, pathways of Zn^2+^ uptake physiologically favor maximal efficient mechanisms, rather than fast channel uptake systems.

Can an H^+^ coupled transporter support greater Zn^2+^ uptake? Assuming a stoichiometry of 5 H^+^ per Zn^2+^ (see Fig. 2C) and based on the Gibbs free energy calculation (see materials and methods), the energy earned from H^+^ transport at pH7.4 would be −0.18674 Kcal/mol, and 17.318 Kcal/mol, at pH5, thus yielding a Zn^2+^ gradient which is 10^6^-fold higher at pH5, than that at a neutral pH. Luminal pH, in the proximal small intestines is in the range of 5.5-7. Thus, our suggested mechanism of acidic pH regulation and H^+^ coupled Zn^2+^ co-transport is also supported by the physiology of the GI tract and would enable higher nutritional Zn^2+^ absorbance, by utilizing the driving force generated by the H^+^ gradient. Correspondingly, the pH gradient across the PM of renal tubules would enable coupling H^+^ and Zn^2+^ transport, to enhance reabsorption of Zn^2+^, and indeed cells of the proximal tubule abundantly express ZIP4 ^26^, as well as other members of the ZIP family ^43^.

Similar mechanisms of enhanced uptake vie either H^+^ or Na^+^ coupling are well documented and abundant in the GI tract ^44^ and renal tubules ^45^. The Na^+^/Glucose co-transporter, for instance, couples Glucose uptake to the Na^+^ gradient across the PM to maximize Glucose absorption from the GI tract, and reabsorption from the proximal tubules of the nephron ^46^. Indeed, in the proximal tubule of the nephron, reabsorption of filtered glucose is 100%. Interestingly, uptake of Glucose from the blood stream into erythrocytes and muscle is mediated by facilitated diffusion that maintains constant basal levels of glucose within cells, per the available glucose gradient. Thus, the comparison of Na^+^/Glucose co-transport and Glucose facilitated diffusion demonstrates in vivo the importance of an optimal mechanism for maximal absorption.

There are, however, risks related to a cellular toxic surge of Zn^2+ 47, 48^ following a bolus of Zn^2+^ in the digestive system. This toxic ionic surge can be encountered by several documented ZIP4 “safety valve mechanisms”; (a) the slow rate of ZIP4 activity limits Zn^2+^ uptake and thus prevents potential toxicity, (b) the transporter undergoes rapid endocytosis ^22, 23^ that limits the duration of Zn^2+^ uptake, and (c) following the uptake of Zn^2+^ from the digestive tract or other absorption tissues, such as the renal tubules, Zn^2+^ can be rapidly transported, vectorially, into vesicular compartments by the activity of vesicular ZnTs ^49^ or across the cells’ plasma membrane via ZnT1 ^50^.

Remarkably, like ZIP4, the renal Na^+^/Glucose co-transporter SGLT1 has also been suggested to undergo a regulatory process of endocytosis ^51^, suggesting these diverse mechanisms did not only develop to support similar energy considerations, but also harbor similar regulatory strategies.

### Genetic variants associated with Zn^2+^ defiantly, in AE patients, are linked to either catalytic or non-catalytic domains of ZIP4

Genetic variation in ZIP4 is linked to Zn^2+^ deficiency in human subjects ^25-28^. Several of these mutations lead to deletion and frame shift mutations, however, the majority result in single amino acid substitutions. Several of these lead to failure of ZIP4 accumulation at the plasma membrane, possibly due to mis-folding that disrupts glycosylation sites and leads to failure in protein localization ^28^.

Two variants that were previously tested (P200L and G539R) differed from the rest, in that they did accumulate at the plasma membrane, yet showed diminished Zn^2+^ uptake over a 15-minute time course assay ^28^. Our results support the finding that ZIP4^P200L^ localizes to the PM, however, under our shorter 2-minute experiment, Zn^2+^ transport rates were no different from those of the wild type protein, indicating that catalytic Zn^2+^ transport was not impaired by this mutation. A possible explanation to this discrepancy could be the location of this residue, at the extra-cellular N-terminal domain of ZIP4 that has been shown to be cleaved under Zn^2+^ deficiency and play part in the regulatory process this protein undergoes ^52^. We propose this mutated transporter is catalytically active, but undergoes different regulation following Zn^2+^ exposure, which would explain the diminished uptake of Zn^2+^, following a 15 minute experiment that allows endocytosis, but not observed by us after 2 minutes. Thus, the apparent reduction in Zn^2+^ transport following the longer time-course may in fact be related to decreased availability of ZIP4 at the PM. This finding is consistent with the recently published 3D-structure of BbZIP, on which the structure of hZIP4 was mapped ^30^.

ZIP4^G539R^ also shows diminished Zn^2+^ uptake over a 15-minute experiment ^52^. Similarly, this mutant also displayed aberration of direct Zn^2+^ transport in our 2 minute transport assay, which given the proximity of this mutation to the residues forming the putative catalytic ion transport core, could be attributed directly to the catalytic activity of ZIP4, by disruption of either Zn^2+^ binding or to the conformational changes the transporter undergoes.

Unlike the wild type transporter, Zn^2+^ does not regulate endocytosis of ZIP4^P200L^ and ZIP4^G539R^. Taken together with diminished Zn^2+^ uptake by ZIP4^P200L^, following a 15-minute assay, but not after a 2-minute assay, these results suggest endocytosis of ZIP4 plays not only a regulatory role, but also plays a yet poorly understood part in Zn^2+^ uptake.

## Materials and Methods

### Cell Cultures

HEK293-T cells (human embryonic kidney cell line) were cultured in Dulbecco’s modified Eagle’s medium (DMEM), supplemented with 10% Fetal Calf Serum (FCS), 1% Streptomycin and 1% Penicillin. Cells were grown in either 25cm^2^ or 75cm^2^ flasks, in a humidified CO_2_ incubator, at 37°C.

For live-cell imaging and immunocytochemistry experiments, cells were transferred on to glass cover slips, in 60mm cell culture dishes. For immunoblotting, cells were transferred to 100-mm cell culture dishes.

### Plasmid transfection

Cells were transfected with 0.67µg of the indicated HA tagged mZIP4 double-stranded plasmid (accession number BC023498) using the well documented CaPO4 precipitation protocol in cultures of 40-60% confluence, 48 hours prior to experiment. The various plasmids used for transfection are described in the following section.

Site-directed mutagenesis was performed using the QuikChange site-directed mutagenesis kit (Stratagene) according to the following protocol:

**Table 3:** Site directed mutagenesis reaction components.

| Component | Amount |
| --- | --- |
| dsDNA template<br>(100ng) | 0.333µl |
| Primer F (0.3µM) | 0.75µl |
| Primer R (0.3µM) | 0.75µl |
| dNTPs | 1.5µl |
| Enzyme | 0.5µl |
| ddH<sub>2</sub>O (a total of 25µl<br>) | 21.167µl |
| Total volume | 25µl |

**Table 4:** Site directed mutagenesis PCR

| Cycles | Step | Temperature | Time |
| --- | --- | --- | --- |
| 1 | Initial denaturation | 95°C | 5:00 |
| 20 | Denaturation | 98°C | 0:20 |
|  | Anealing | 78°C | 0:20 |
|  | Elongation | 72°C | 0:30/1kb template |
| 1 | Final elongation | 78°C | 5:00 |

The following primers were designed using the primer design tool, on the University of Washington server (http://depts.washington.edu/bakerpg/webtools/PD.html) and manufactured by SIGMA.

ZIP4^P200L^: CCAAGGCCTG<u>**CTT**</u>AGCCCTCAGTA.

ZIP4^G539R^: CCATGAACTC<u>**CGC**</u>GACTTCGCTGCTCTG

### Immunoblot analysis

Cells were extracted using 200µl of boiling denaturative lysis buffer (1% SDS, 10mM Tris-HCl, pH8) / 100-mm plate and transferred to ice. A protease inhibitor mixture (Boehringer Complete protease inhibitor mixture; Roche Applied Science) was added to the lysates, and protein concentrations were determined using the modified Lowry procedure ^53^. SDS-PAGE and immunoblot analyses were performed, using anti-actin and anti-HA antibodies at dilutions of 1:40000, and 1:2000 respectively. Secondary anti-mouse and anti-rabbit antibodies (Jackson Immunoesearch) were used at dilutions of 1:20000 and 1:40000, respectively.

The detection of membrane embedded ZIP4 the protocol described by Kim et al ^22^ was used. Specifically, cells were incubated with 50µM Zn^2+^ for 0-60 minutes, as indicated. Zn^2+^ uptake and endocytosis were terminated by transferring the cells to ice and subsequently washing the cells with ice-cold PBS. Following this step, cells were fixed using 4% PFA in 0.1M PBS. Cells were then washed in PBS to remove residual PFA, and following the removal of PBS, cells were incubated with 1µg/µl anti-HA antibody, for 30 minutes at room temperature. Cells were then washed 5 times with PBS, to remove residual unbound antibodies and immediately exposed to boiling denaturative lysis buffer for western blotting. Following SDS-PAGE and immunobloting, membranes were incubated with secondary anti-mouse antibody.

### Imaging system

The imaging system consisted of an Axiovert 100 inverted microscope (Zeiss), Polychrome II monochromator (TILL Photonics, Planegg, Germany) and a SensiCam cooled charge-coupled device (PCO, Kelheim, Germany). Fluorescent imaging measurements were acquired with the Imaging Workbench 6 software (Axon Instruments, Foster City, CA) and analyzed using Microsoft exel, Kaleidograph and Matlab.

### Live cell fluorescent imaging

Cytoplasmic Zn^2+^ transport was determined in cells loaded with 0.5µM Fluozin-3AM. To verify that the fluorescence changes were related to intracellular ions, the cell-permeable heavy metal chelator *N*,*N*,*N*’,*N’*-tetrakis-(2-pyridylmethyl)-ethylenediamine (TPEN; 20 µM) was used.

Cytoplasmic pH changes, indicative of H^+^ transport, were determined in cells loaded with 1 µM BCECF-AM, a pH sensitive dye. Results were calibrated using high K^+^ Ringer’s solution set to pH values of 6–8 in the presence of Nigericin ^54^.

Intracellular acidification was triggered using the ammonium prepulse paradigm ^41^. Cells were superfused with Ringer’s solution containing NH4Cl (30mM, replacing the equivalent 30mM NaCl), which was subsequently replaced with NH4Cl-free solution, thus triggering intracellular acidification.

### Statistics

Data analysis was performed using the SPSS software (version 14.0; SPSS Inc., Chicago, IL). All results shown are the means ± S.E. of at least three individual experiments (*n* ≥ 3). *t* test *p* values of ≤ 0.05 were considered significant following Levene’s test for equality of variances. Significance of the results is as follows: relative to control (*) or relative to ZnT5 / ZIP4 (#). *p* ≤ 0.05 for all experiments, unless otherwise stated.

## References

1 Vallee, B.L. & Falchuk, K.H. The biochemical basis of zinc physiology. Physiol-Rev 73, 79–118 (1993).

2 Prasad, A.S. Zinc deficiency. BMJ 326, 409–410 (2003).

3 Andreini, C., Banci, L., Bertini, I. & Rosato, A. Counting the zinc-proteins encoded in the human genome. J Proteome Res 5, 196–201 (2006).

4 Eide, D.J. Zinc transporters and the cellular trafficking of zinc. Biochim Biophys Acta. 1763, 711–722 (2006).

5 Chimienti, F., Devergnas, S., Favier, A. & Seve, M. Identification and cloning of a beta-cell-specific zinc transporter, ZnT-8, localized into insulin secretory granules. Diabetes 53, 2330–2337. (2004).

6 Chimienti, F. et al. In vivo expression and functional characterization of the zinc transporter ZnT8 in glucose-induced insulin secretion. J. Cell. Sci. 119, 4199–4206 (2006).

7 Suzuki, T. et al. Two different zinc transport complexes of cation diffusion facilitator proteins localized in the secretory pathway operate to activate alkaline phosphatases in vertebrate cells. J Biol Chem 280, 30956–30962 (2005).

8 Suzuki, T. et al. Zinc transporters, ZnT5 and ZnT7, are required for the activation of alkaline phosphatases, zinc-requiring enzymes that are glycosylphosphatidylinositol-anchored to the cytoplasmic membrane. J. Biol. Chem. 280, 637–643 (2005).

9 Ohana, E. et al. Identification of the Zn2+ binding site and mode of operation of a mammalian Zn2+ transporter. J Biol Chem 284, 17677–17686 (2009).

10 Kawachi, M., Kobae, Y., Mimura, T. & Maeshima, M. Deletion of a histidine-rich loop of AtMTP1, a vacuolar Zn(2+)/H(+) antiporter of Arabidopsis thaliana, stimulates the transport activity. J Biol Chem 283, 8374–8383 (2008).

11 Chao, Y. & Fu, D. Kinetic study of the antiport mechanism of an Escherichia coli zinc transporter, ZitB. J Biol Chem. 279, 12043–12050. (2004).

12 Lu, M. & Fu, D. Structure of the zinc transporter YiiP. Science 317, 1746–1748 (2007).

13 Jeong, J. et al. Promotion of vesicular zinc efflux by ZIP13 and its implications for spondylocheiro dysplastic Ehlers-Danlos syndrome. Proc Natl Acad Sci U S A 109, E3530–3538 (2012).

14 Giunta, C. et al. Spondylocheiro dysplastic form of the Ehlers-Danlos syndrome--an autosomal-recessive entity caused by mutations in the zinc transporter gene SLC39A13. Am J Hum Genet 82, 1290–1305 (2008).

15 Fukada, T. et al. The zinc transporter SLC39A13/ZIP13 is required for connective tissue development; its involvement in BMP/TGF-beta signaling pathways. PLoS ONE 3, e3642 (2008).

16 Dalton, T.P. et al. Identification of mouse SLC39A8 as the transporter responsible for cadmium-induced toxicity in the testis. Proc. Natl. Acad. Sci. USA 102, 3401–3406. Epub 2005 Feb 3418. (2005).

17 Gitan, R.S., Shababi, M., Kramer, M. & Eide, D.J. A cytosolic domain of the yeast Zrt1 zinc transporter is required for its post-translational inactivation in response to zinc and cadmium. J Biol Chem 278, 39558–39564 (2003).

18 Taylor, K.M. & Nicholson, R.I. The LZT proteins; the LIV-1 subfamily of zinc transporters. Biochim. Biophys. Acta 1611, 16–30. (2003).

19 Rogers, E.E., Eide, D.J. & Guerinot, M.L. Altered selectivity in an Arabidopsis metal transporter. Proc Natl Acad Sci U S A 97, 12356–12360 (2000).

20 Hoch, E. et al. Histidine pairing at the metal transport site of mammalian ZnT transporters controls Zn2+ over Cd2+ selectivity. Proc Natl Acad Sci U S A 109, 7202–7207 (2012).

21 Dufner-Beattie, J. et al. The acrodermatitis enteropathica gene ZIP4 encodes a tissue-specific, zinc-regulated zinc transporter in mice. J Biol Chem 278, 33474–33481 (2003).

22 Kim, B.E. et al. Zn2+-stimulated endocytosis of the mZIP4 zinc transporter regulates its location at the plasma membrane. J. Biol. Chem. 279, 4523–4530 (2004).

23 Weaver, B.P., Dufner-Beattie, J., Kambe, T. & Andrews, G.K. Novel zinc-responsive post-transcriptional mechanisms reciprocally regulate expression of the mouse Slc39a4 and Slc39a5 zinc transporters (Zip4 and Zip5). Biol Chem 388, 1301–1312 (2007).

24 Dufner-Beattie, J., Kuo, Y.M., Gitschier, J. & Andrews, G.K. The adaptive response to dietary zinc in mice involves the differential cellular localization and zinc regulation of the zinc transporters ZIP4 and ZIP5. J Biol Chem 279, 49082–49090. Epub 42004 Sep 49089. (2004).

25 Kury, S. et al. Identification of SLC39A4, a gene involved in acrodermatitis enteropathica. Nat Genet 31, 239–240. Epub 2002 Jun 2017. (x2002).

26 Wang, K., Zhou, B., Kuo, Y.M., Zemansky, J. & Gitschier, J. A novel member of a zinc transporter family is defective in acrodermatitis enteropathica. Am J Hum Genet 71, 66–73. Epub 2002 May 2024. (2002).

27 Nakano, A., Nakano, H., Nomura, K., Toyomaki, Y. & Hanada, K. Novel SLC39A4 mutations in acrodermatitis enteropathica. J Invest Dermatol 120, 963–966 (2003).

28 Wang, F. et al. Acrodermatitis enteropathica mutations affect transport activity, localization and zinc-responsive trafficking of the mouse ZIP4 zinc transporter. Hum Mol Genet 13, 563–571 (2004).

29 Prasad, A.S. The role of zinc in gastrointestinal and liver disease. Clin Gastroenterol 12, 713–741 (1983).

30 Zhang, T. et al. Crystal structures of a ZIP zinc transporter reveal a binuclear metal center in the transport pathway. Sci Adv 3, e1700344 (2017).

31 Li, M. et al. Aberrant expression of zinc transporter ZIP4 (SLC39A4) significantly contributes to human pancreatic cancer pathogenesis and progression. Proc Natl Acad Sci U S A 104, 18636–18641 (2007).

32 Li, M. et al. Down-regulation of ZIP4 by RNA interference inhibits pancreatic cancer growth and increases the survival of nude mice with pancreatic cancer xenografts. Clin Cancer Res 15, 5993–6001 (2009).

33 Gaither, L.A. & Eide, D.J. Functional expression of the human hZIP2 zinc transporter. J Biol Chem 275, 5560–5564 (2000).

34 He, L. et al. ZIP8, member of the solute-carrier-39 (SLC39) metal-transporter family: characterization of transporter properties. Mol Pharmacol 70, 171–180 (2006).

35 Girijashanker, K. et al. Slc39a14 gene encodes ZIP14, a metal/bicarbonate symporter: similarities to the ZIP8 transporter. Mol Pharmacol 73, 1413–1423 (2008).

36 Qiu, A. & Hogstrand, C. Functional expression of a low-affinity zinc uptake transporter (FrZIP2) from pufferfish (Takifugu rubripes) in MDCK cells. Biochem J 390, 777–786 (2005).

37 Antala, S. & Dempski, R.E. The human ZIP4 transporter has two distinct binding affinities and mediates transport of multiple transition metals. Biochemistry 51, 963–973 (2012).

38 Lin, W., Chai, J., Love, J. & Fu, D. Selective electrodiffusion of zinc ions in a Zrt-, Irt-like protein, ZIPB. J Biol Chem 285, 39013–39020 (2010).

39 Kiedrowski, L. Cytosolic acidification and intracellular zinc release in hippocampal neurons. Journal of neurochemistry 121, 438–450 (2012).

40 Sensi, S.L. et al. Modulation of mitochondrial function by endogenous Zn2+ pools. Proc. Natl. Acad. Sci. USA 100, 6157–6162. (2003).

41 Azriel-Tamir, H., Sharir, H., Schwartz, B. & Hershfinkel, M. Extracellular zinc triggers ERK-dependent activation of Na+/H+ exchange in colonocytes mediated by the zinc-sensing receptor. J. Biol. Chem. 279, 51804–51816. (2004).

42 Outten, C.E. & O’Halloran, T.V. Femtomolar Sensitivity of Metalloregulatory Proteins Controlling Zinc Homeostasis. Science 292, 2488–2492 (2001).

43 Wang, F. et al. Zinc-stimulated endocytosis controls activity of the mouse ZIP1 and ZIP3 zinc uptake transporters. J. Biol. Chem. 279, 24631–24639 (2004).

44 Thwaites, D.T. & Anderson, C.M. H+-coupled nutrient, micronutrient and drug transporters in the mammalian small intestine. Exp Physiol 92, 603–619 (2007).

45 Hummel, C.S. et al. Glucose transport by human renal Na+/D-glucose cotransporters SGLT1 and SGLT2. Am J Physiol Cell Physiol 300, C14–21 (2011).

46 Wright, E.M. Renal Na(+)-glucose cotransporters. Am J Physiol Renal Physiol 280, F10–18 (2001).

47 Choi, D.W. & Koh, J.Y. Zinc and brain injury. Annu. Rev. Neurosci. 21, 347–375 (1998).

48 Weiss, J.H., Sensi, S.L. & Koh, J.Y. Zn(2+): a novel ionic mediator of neural injury in brain disease. Trends Pharmacol. Sci. 21, 395–401. (2000).

49 Kambe, T. Molecular architecture and function of ZnT transporters. Curr Top Membr 69, 199–220 (2012).

50 Qin, Y., Thomas, D., Fontaine, C.P. & Colvin, R.A. Silencing of ZnT1 reduces Zn2+ efflux in cultured cortical neurons. Neurosci Lett 450, 206–210 (2009).

51 Wright, E.M., Hirsch, J.R., Loo, D.D. & Zampighi, G.A. Regulation of Na+/glucose cotransporters. J Exp Biol 200, 287–293 (1997).

52 Kambe, T. & Andrews, G.K. Novel proteolytic processing of the ectodomain of the zinc transporter ZIP4 (SLC39A4) during zinc deficiency is inhibited by acrodermatitis enteropathica mutations. Mol. Cell. Biol. 29, 129–139. (2009).

53 Markwell, M.A., Haas, S.M., Bieber, L.L. & Tolbert, N.E. A modification of the Lowry procedure to simplify protein determination in membrane and lipoprotein samples. Anal Biochem 87, 206–210 (1978).

54 Machen, T.E. et al. pH of TGN and recycling endosomes of H+/K+-ATPase-transfected HEK-293 cells: implications for pH regulation in the secretory pathway. Am. J. Physiol. Cell Physiol. 285, C205–214 (2003).

